## Supplementary Figures for "Foxp3 Orchestrates Reorganization of Chromatin Architecture to Establish Regulatory T Cell Identity"

A

Gating strategy for T cells in the thymus

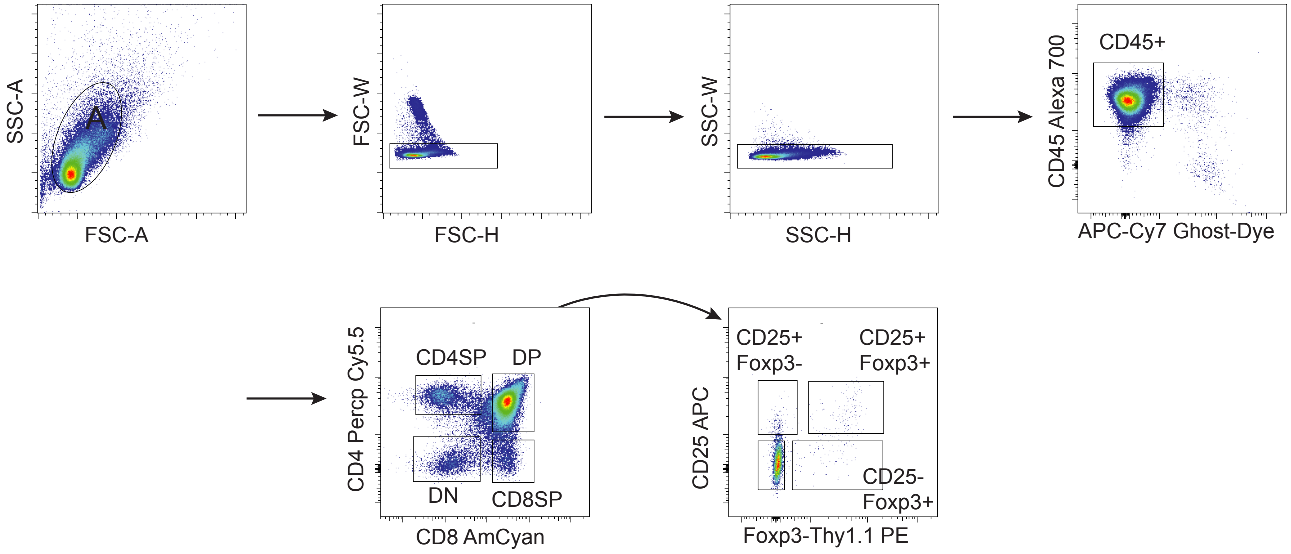


B

Gating strategy for Treg and Tcon cells in the spleen

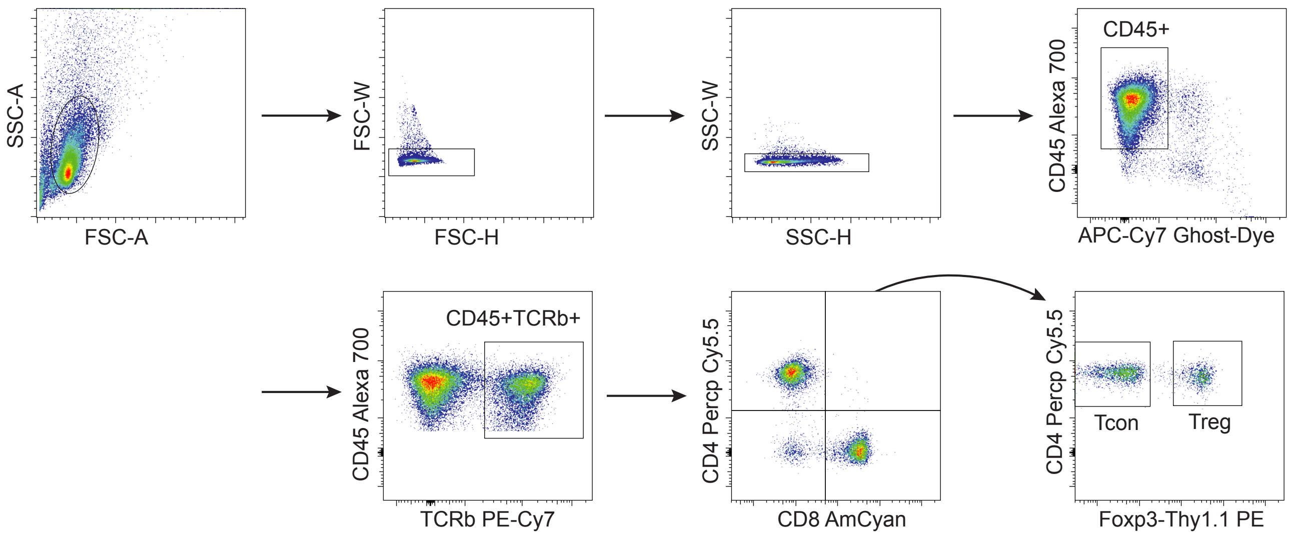


**Supplementary Figure 1**

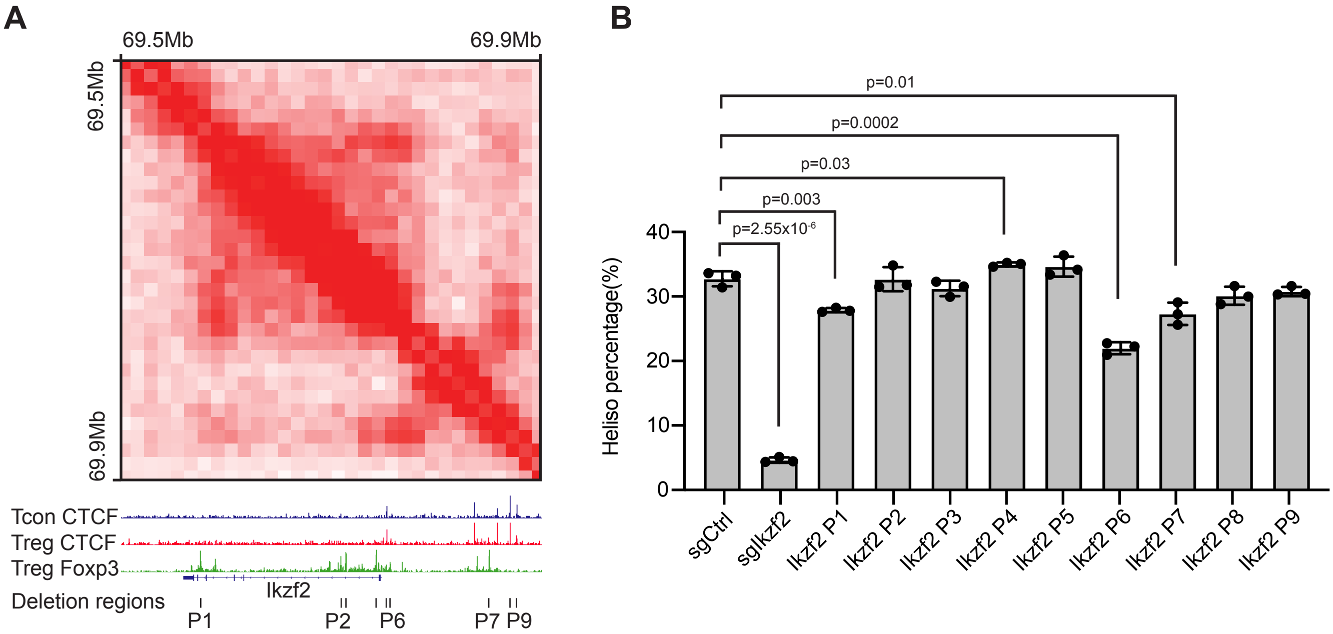


**Supplementary Figure 2**

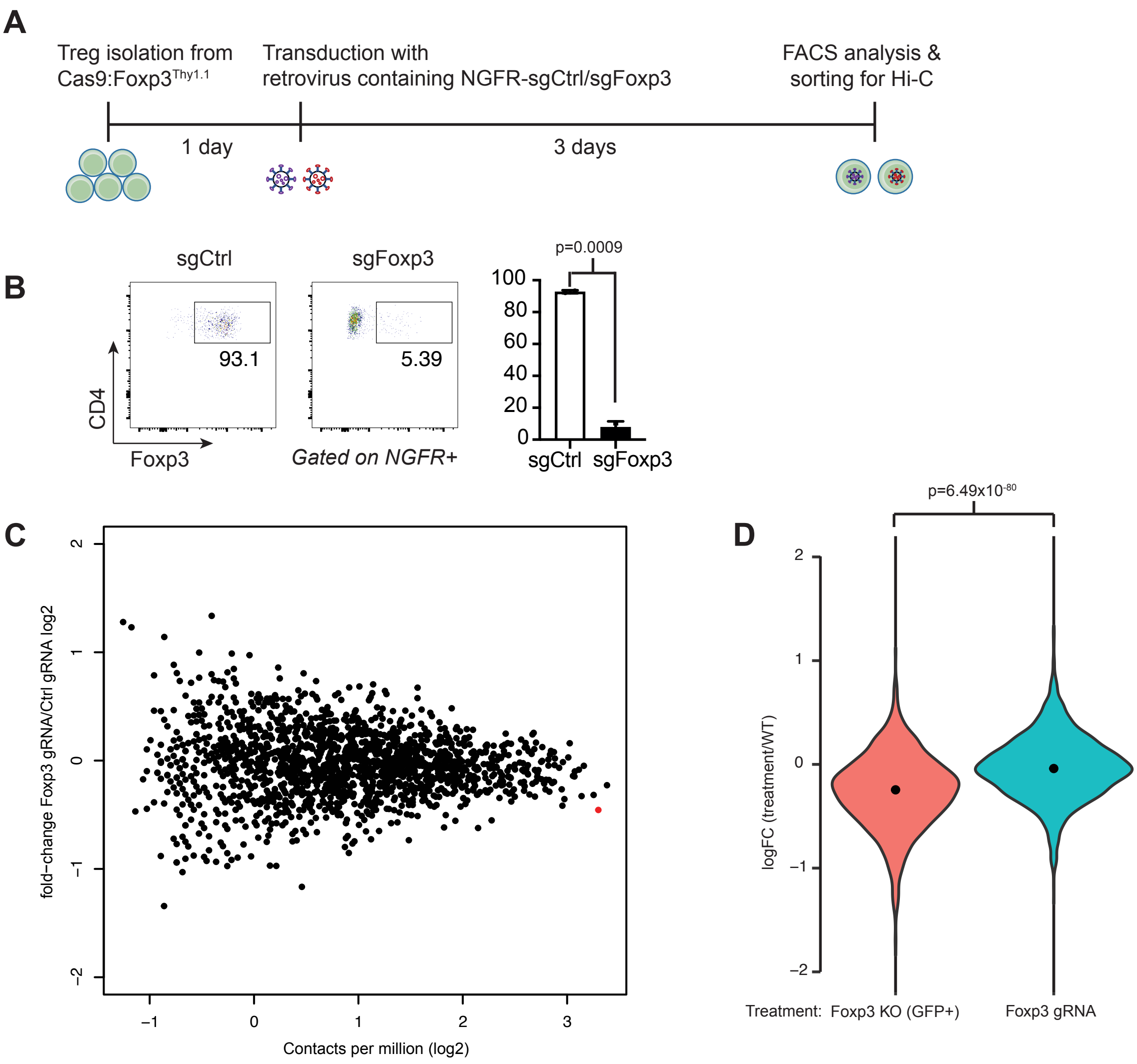


**Supplementary Figure 3**
